# The Human Motor Cortex Contributes to Gravity Compensation to Maintain Posture and During Reaching

**DOI:** 10.1101/723056

**Authors:** R. L. Hardesty, P. H. Ellaway, V. Gritsenko

**Author notes:** **Address for Correspondence** V. Gritsenko, PO Box 9226, One Medical Center Drive, Morgantown, WV, 26505-9226.

## Abstract

How the neural motor system recruits muscles to support the arm against gravity is a matter of active debate. It is unknown how the neural motor system compensates for the changing gravity-related joint moments either when holding a steady-state posture or during movement between postural steady states, e.g., during reaching. Here we used single-pulse transcranial magnetic stimulation to compare the roles that the human primary motor cortex plays in the muscle recruitment to compensate for gravity. We hypothesized that the motor cortex contributes to muscle recruitment to both maintain posture and to compensate for changes in gravitational passive joint moments during movement. To test this hypothesis, we used visual targets in virtual reality to instruct five postures and three movements with or against gravity. We then measured the amplitude and gain of motor evoked potentials in multiple muscles of the arm at several phases of the reaching motion and during posture maintenance. Stimulation below the resting motor threshold, calibrated to the biceps muscle, caused motor evoked potentials in all muscles during all postural and reaching tasks. The amplitude of motor evoked potentials was proportional to the motoneuronal excitability measured as muscle activity. The coefficient of proportionality was positively correlated with the postural component of muscle moment during posture and movement. Altogether our results support the hypothesis. The observed contribution of the motor cortex to the recruitment of multiple antagonistic muscles suggests a whole-limb strategy for overcoming passive gravity-related moments with both active muscle moments and muscle co-activation that modulates limb impedance.

**New & Noteworthy:** Maintaining static posture and producing motion appear to be contradictory tasks for the nervous system. In contrast to this seeming dichotomy, our results show that the motorcortical control signals play the same role in both tasks when it is framed in biomechanical terms, i.e., muscle contractions needed to compensate for gravity.

## Introduction

Posture and movement control are distinct tasks for the neural motor system. Posture requires maintaining a steady state, such as holding the elbow flexed against gravity when typing. In contrast, movement can be viewed as a transition between steady states. Traditionally, the neural control of these tasks has also been subdivided into a focal movement and postural adjustments. An example of the neural command for a focal movement would be the activation of arm muscles to produce a reaching movement, while the postural adjustment would be the activation of trunk muscles that compensate for the change in the body center of mass (Massion 1992). Biomechanically, both movement and posture, including postural adjustments, can be accomplished in two ways, 1) generating active joint moments with muscle contractions, termed muscle torques and 2) changing joint stiffness and viscosity components of mechanical impedance through muscle co-contraction. The forces generated by muscle contraction sum around a joint’s degrees of freedom (DoFs) with directions defined by muscle geometry and moment arms. When they create counterbalanced muscle moments that sum up to zero, that defines the stiffness and viscosity components of mechanical impedance (Zajac 1989). Thus, increasing mechanical impedance through the co-contraction of antagonistic muscles can help maintain posture in presence of internal and external perturbations. However, when the muscle forces do not perfectly balance out around the joint, muscle torques arise. When these muscle torques are equal and opposite to the passive torques caused by gravity and any other external or internal force, no motion is produced or the same posture is maintained. Alternatively, when the amplitude of these muscle torques are larger than the passive torques, then active motion happens. Active motion is under the direct control of the nervous motor system through muscle contractions, we termed this component of motor control focal movement. Alternatively, when the amplitude of muscle torques are smaller than the passive torques, then passive motion results, which is not under the direct control of the nervous system. In this case, modulating mechanical impedance can enable the nervous system to keep control over the passive motion of the limb. Based on these biomechanical considerations, several models of the neural mechanisms for the control of posture and movement have been proposed. These models consist of two parallel or serialized controllers, one for generating the focal movement command through active muscle torques and one for posture maintenance through mechanical impedance (Ghez et al. 2007; Lametti et al. 2007; Todorov and Jordan 2002; Yadav and Sainburg 2011).

In this study of reaching, we further subdivide focal movement into posture- and movement-related components. The posture-related component of focal movement is the transition from the starting posture to a new posture. It is accompanied by the changing gravitational moments at arm joints caused by the different orientation of arm segments relative to the gravity vector. The second movement-related component is the residual muscle activity responsible for propulsion, i.e., acceleration and deceleration phases of motion. These posture- and movement-related components of focal movement can be observed in electromyographic (EMG) profiles. The former corresponds to the static component of EMG defined as the change in EMG amplitude between the starting and ending posture with a linear or cosine transition during movement; the latter is the phasic component of EMG we typically observe as bursts (Buneo et al. 1994; Flanders et al. 1996; Olesh et al. 2017). Here we suggest that the posture-related component of focal movement or the corresponding static EMG is generated by the same neural mechanism as that responsible for posture maintenance, specifically that for postural adjustment during focal movement. To test this idea, we hypothesize that the motor cortex contributes to muscle recruitment to both maintain posture and to compensate for changes in gravitational passive joint moments during movement.

Evidence in cats suggest that postural adjustments are initiated and shaped by the output of the primary motor cortex via the corticospinal tract through its collaterals to the reticulospinal tract (Schepens and Drew 2004; Yakovenko and Drew 2009). In humans, the excitability of the corticospinal tract and its collaterals has been studied using single-pulse transcranial magnetic stimulation (TMS). The descending tracts converge on the motoneurons, the final common pathway. Consequently, TMS over the motor cortex activated the corticospinal tract and its collaterals, which in turn activate spinal motoneurons causing motor evoked potentials (MEPs) that can be observed in EMG. MEPs provide a measure of instantaneous excitability or recruitment of both pre- and post-synaptic elements of the corticospinal tract and its collaterals (Bestmann and Krakauer 2015; Di Lazzaro et al. 2004; Reis et al. 2008; Rothwell 1997). TMS at low intensity does not disrupt neural state (Cros et al. 2007), therefore changes in MEP amplitudes can reveal the changes in the descending contribution to muscle recruitment during posture and movement. For example, MEP amplitudes has been shown to change with different hand postures reflecting different corticospinal contribution to the recruitment of muscles supporting those postures (Perez and Rothwell 2015). The amplitude of a given MEP also depends on the excitability of the motoneuron pool of the parent muscle (Darling et al. 2006; Groppa et al. 2012; MacKinnon and Rothwell 2000; Taylor 2006). However, the proportional contribution of the presynaptic elements to muscle recruitment, termed corticospinal gain, can be estimated as the proportion between MEP amplitude and the amplitude of the recruitment of motoneuronal pools observed with EMG in absence of MEPs (Gritsenko et al. 2011). During movement the corticospinal gain is modulated in a phase-dependent manner independently from the profile of EMG in wrist muscles and proximal antigravity muscles (Lemon et al. 1995). Here we directly compare the corticospinal gain in the same muscles involved in both posture and movement to determine if the changes in corticospinal gain reflect the changes in gravity load during both posture and movement.

## Materials and Methods

### Subjects

The Institutional Review Board approved all procedures in this study (Protocol #1309092800). Potential participants with any musculoskeletal pathologies or injuries, prior history of seizures, fainting, or tinnitus, or those taking prescribed psychoactive medications were excluded. We obtained informed consent prior to the start of experiments. We recruited 10 healthy individuals (6 male, 4 female, 24.3 ± 1.8 years old, 76.3 ± 14.5 kg). All participants reported to be right-hand dominant.

### Motion Capture and Tasks

During the experiment, arm postures and reaching goals were defined in virtual reality environment created using Vizard software (WorldViz) and Oculus headset (Fig. 1A). Spherical virtual targets 8 cm in diameter were displayed in a sagittal plane and defined a set of postures and reaching tasks as described in detail in (Hardesty et al. 2020) (Fig. 1B). The movements were similar to the planar movements in a horizontal plane in Gritsenko et al. (2011) but occurred in a vertical sagittal plane without external mechanical constraints.

**Figure 1.**
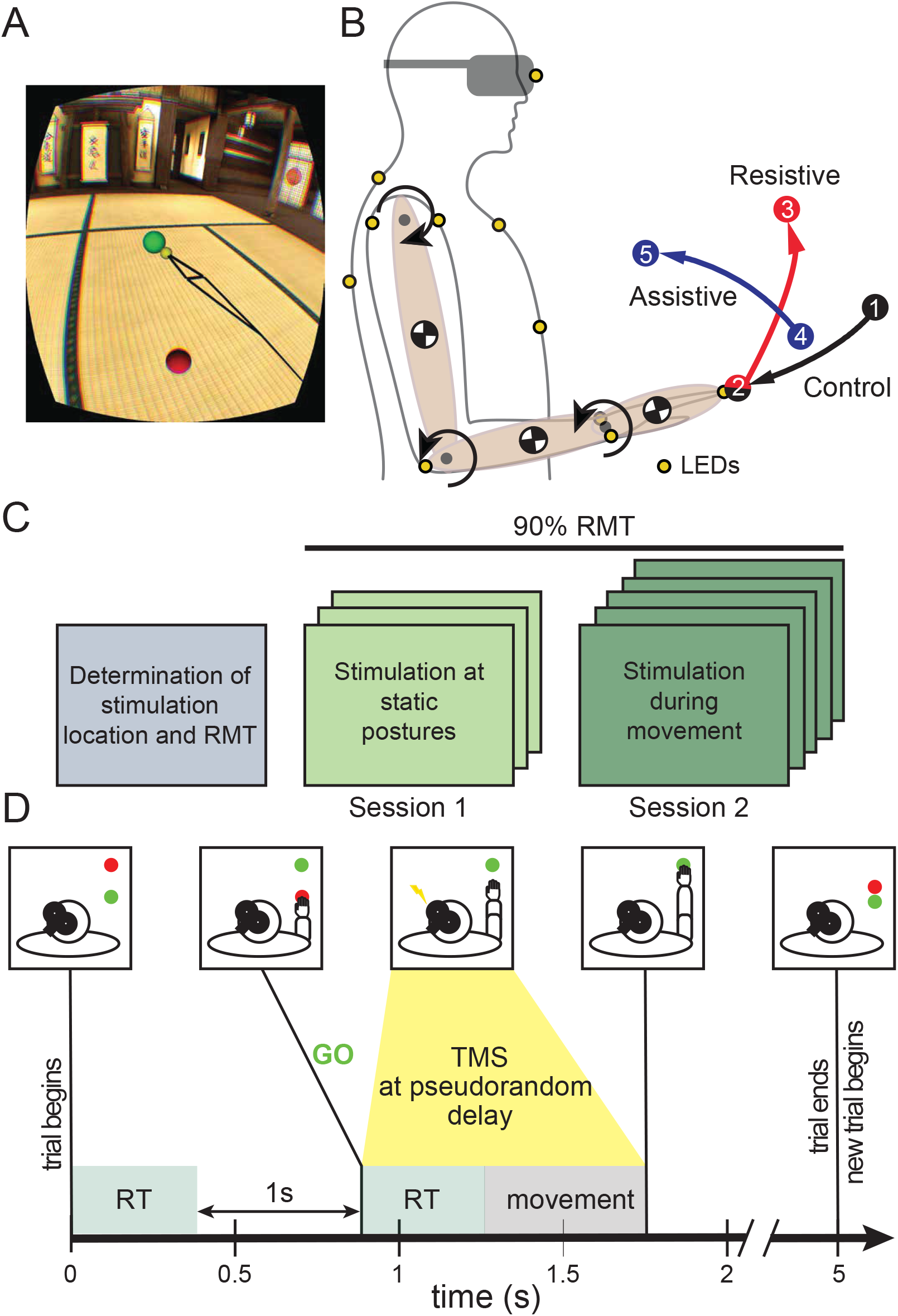
Experimental paradigm. A. Participant’s view of the tasks in virtual reality. Black lines connect virtual representation of LED markers and show the arm position and orientation in virtual reality relative to targets. The yellow sphere represents the fingertip LED on the index finger, which the participants were instructed to move to the center of the green target. B. Schematic of the dynamic model used to calculate joint angles and torques from motion capture and representative endpoint trajectories from the three tasks. Three segments (pink ovals) were used to simulate inertial properties of upper arm, forearm, and hand; centers of mass are indicated by black and white circles. Positive directions for each joint (DOF) are shown with black circular arrows. The start and stop target locations are shown for the Control, Resistive, and Assistive tasks (black, red, and blue, respectively). The motion capture marker locations are shown as yellow circles. C. Visual representation of the experimental paradigm. D. Example timeline of events during a single trial of a movement task. RT stands for reaction time; GO represents the color cue to start the movement.

To minimize the inter-subject variability in kinematics during experiments, the locations of virtual targets were adjusted based on individual segment lengths. The targets were displayed relative to each individual’s shoulder marker on the acromion (Fig. 1B). The experiment was subdivided into two sessions, Session 1 with static postures and Session 2 with reaching movements. In both sessions the participants were asked to reach to or hold at the virtual target with their right (dominant) arm without moving the trunk while keeping the elbow close to the trunk and a neutral, pronated wrist.

During Session 1, participants were asked to reach to a virtual target and hold there for up to 1 minute before moving to the next target and holding there. The starting position for the Resistive task was the same as the ending position for the Control task, therefore 5 postures were explored (Fig. 1B). While the participant was holding the arm in one posture, we applied 12 TMS pulses more than 5s apart at 90% of resting motor threshold (RMT, see below for details). Six or more TMS pulses has been shown to provide sufficiently reliable estimates of MEPs (Lewis et al. 2014).

During Session 2, each trial started with the appearance of start (green) and stop (red) targets (Fig. 1C). The colors and target locations did not change until the participants placed their index finger, indicated with a yellow sphere, into the start target. One second after this occurred, the stop target changed color from red to green, directing individuals to begin the movement (Fig. 1C). The target pairs for a given task were presented to each individual in the same pseudorandom order generated prior to the study. Each task was repeated 138 times (total of 414 trials). The movement trials were divided into stimulation trials (126 per movement, 378 total) and non-stimulation control trials (12 per movement, 36 total) interspersed randomly. During the stimulation trials TMS was performed at 90% of the RMT. In one half of trials (189 of 378), the TMS was triggered once per trial at a random delay of 0 – 550 ms after the participant touched the start target. This triggering method ensured that MEPs were observed either preceding or around the movement onset time. In the other half of stimulation trials, the TMS pulses were triggered once per trial after the participant left the start target at a random delay of 0 – 550 ms. This triggering method ensured that MEPs were observed through the duration of the movement. The timing of each TMS pulse was recorded for post-hoc normalization to the movement phase and binning described below.

Motion capture was performed using the Impulse system (PhaseSpace). Nine light-emitting diode markers were placed on bony landmarks of the arm and trunk using the best practice guidelines (Robertson et al. 2013). Marker coordinates were sampled at 480 Hz using Recap software (PhaseSpace). The marker locations were also streamed to virtual reality and used to represent the three main segments of the arm (hand, forearm, and upper arm) as a stick figure (Fig. 1A). Motion capture, electromyography (EMG, details below), and virtual events were synchronized using custom hardware as described in (Talkington et al. 2015).

### Joint torques

After experiments, motion capture data from Session 2 were low pass filtered at a cutoff frequency of 10Hz. The mean residuals of marker triangulation ± standard deviation across individuals were 5.7 ± 0.45 mm. Joint angles were calculated by defining local coordinate systems of four rigid bodies for the trunk, humerus, forearm, and hand using at least 3 markers per rigid body (Fig. 1B). To calculate active joint torques, we used a dynamical model similar to that used for the selection of tasks as described in (Hardesty et al. 2020). The model was customized to individual morphology by scaling segment length and inertia based on individual’s height and weight using published average proportions (Winter 2009). Inverse simulations were run with the scaled models using the angular kinematic trajectories averaged per task to obtain applied joint torques for a given individual and task, termed muscle torques. The muscle torques were verified using forward simulations and the simulated angular kinematics was compared to the angular kinematics obtained from motion capture data by calculating a root-mean-squared error (RMSE: 0.059 ± 0.035 rad). Positive rotations defined by local coordinate systems are illustrated in Fig. 1B. Analysis was done on the angles and torques around the X axes, i. e. flexion/extension degrees of freedom at all three joints, because motion was primarily in the sagittal plane and out-of-plane torques were negligible. The postural component of muscle torque, termed postural torque, was extracted by running simulations without gravity and subtracting those from the muscle torques simulated in presence of gravity as described in detail in (Olesh et al. 2017). The postural torque has been shown to follow closely the static component of EMG during similar reaching movements (Olesh et al. 2017).

### Electromyography

Muscle activity and responses to TMS were recorded in twelve upper limb muscles using Trigno (Delsys Inc.), a wireless surface EMG system. The recorded muscles included four muscles spanning the shoulder, three muscles spanning both shoulder and elbow, two muscles spanning only the elbow, and three muscles spanning the wrist (Table 1). Muscles were identified based on anatomical landmarks and palpation during contraction; EMG sensors were placed on muscle bellies oriented longitudinally along the muscle fibers EMG signals were sampled at 2 kHz with a gain of 1000. EMG recordings were high pass filtered at 10Hz to remove any signal drift and rectified prior to any EMG/MEP quantification. EMG profiles from trials without TMS were low pass filtered at a cutoff frequency of 20Hz and normalized per individual using a maximum value for each muscle across all tasks.

**Table 1:**
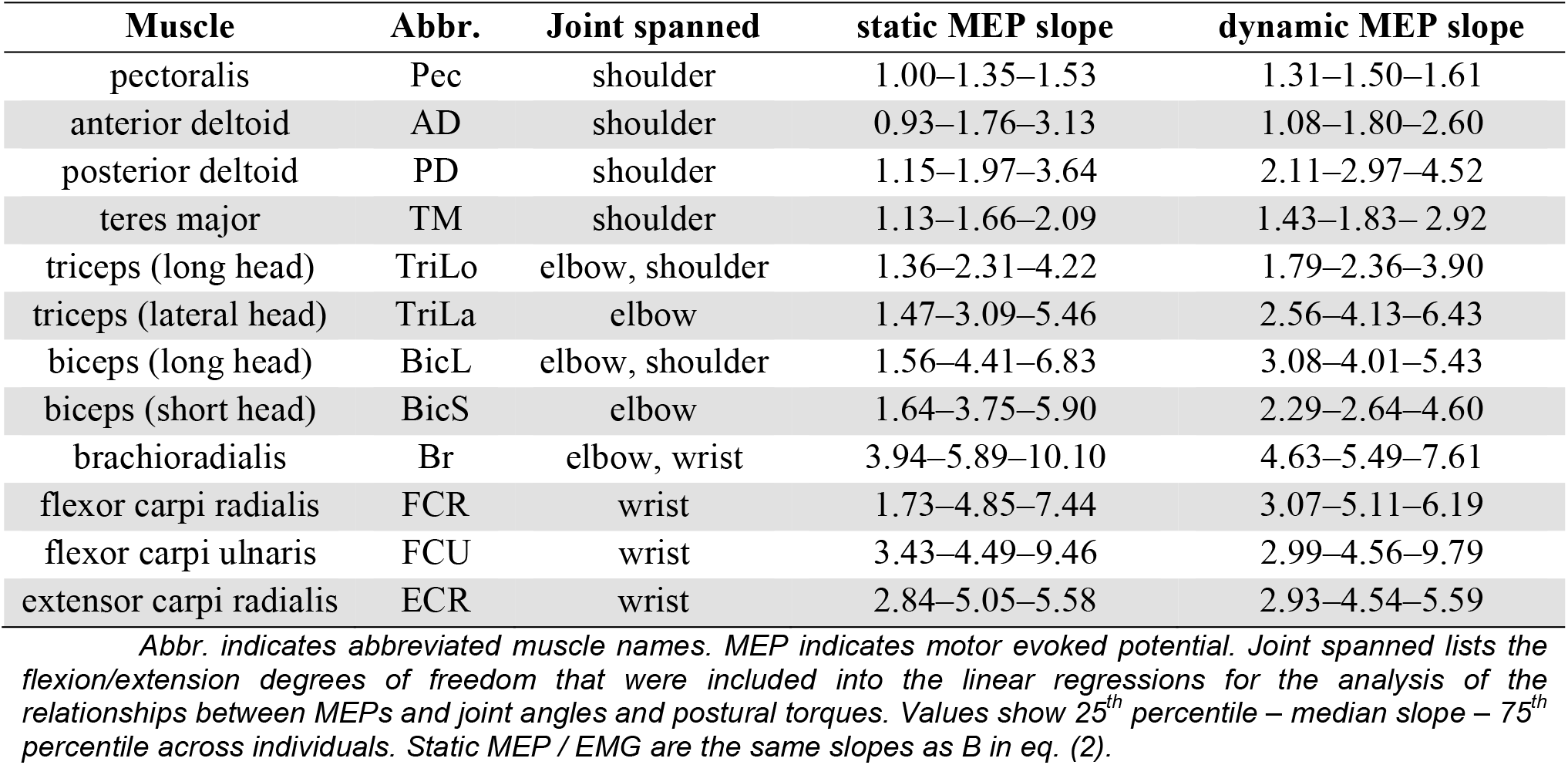
Muscle nomenclature and MEP scaling with background EMG.

### Transcranial magnetic stimulation

We assessed the corticospinal contribution to individual muscles using single-pulse TMS delivered by the Super Rapid stimulator with a figure-of-eight coil (Magstim). The coil was oriented tangentially to the scalp, oriented at a 45° angle to the midline with the handle pointing posteriorly and laterally (Fig. 1C). The coil location over the scalp and its orientation was maintained using the Brainsight neuronavigation system (Rogue Research). This coil positioning has been shown to preferentially activate pyramidal tract neurons (Kaneko et al. 1996; Di Lazzaro et al. 1998; Nakamura et al. 1996) and to generate responses proportional to cortical activity

Focal TMS with the figure-of-eight coil causes MEPs in multiple muscles (Roth et al. 1991; Tofts 1990). The amplitudes of these MEPs and which muscles are involved depends on the strength of the magnetic field generated by the TMS device, the location and orientation of the coil over the scalp, and individual anatomy. The location of the coil was selected using the hot-spot method (Ellaway et al. 1998; Traversa et al. 1997), during which the coil was moved over the estimated location of the primary motor cortex until a location with at least 50 μV motor evoked potential (MEP) in biceps was evoked. This controlled for the anatomical differences between individuals and defined a consistent stimulation location on the motor homunculus (Penfield and Rasmussen 1950). This was done with the anticipation that stimulating the same anatomical location with the lowest corticospinal excitability measured in biceps would activate the same component of the whole corticospinal tract across individuals and produce similar responses in other muscles across individuals.

The strength of the magnetic field is typically tailored to individuals using a resting motor threshold (RMT) method, i.e., identifying the lowest magnetic field strength that evokes MEPs 50% of the time. This reduces the inter-subject variability in TMS responses. RMT was determined at the hot-spot location by varying the stimulation intensity until a MEP > 50 μV was evoked 50% of the time in BicL. This procedure ensured that the stimulation amplitude was adjusted to individual differences in corticospinal excitability at rest at the time of experiment and thus, further minimized inter-subject differences in MEP amplitudes. A single TMS pulse was delivered in a given trial and each trial lasted 5 s. This was done to minimize the instances when two TMS pulses would be delivered at > 0.2 Hz instantaneous frequency to avoid any long-term changes in corticospinal excitability (Chen et al. 1997). The experiment consisted of two consecutive sessions conducted on the same day. The number of repetitions of some conditions was limited due to time constraints in efforts of preventing fatigue.

The amplitudes of MEPs produced using the described method are stable across days (Pascual-Leone et al. 1995; Uy et al. 2002) and have sufficient spatial resolution to differentiate proximal and distal muscles (Pascual-Leone et al. 1997). Here MEPs were quantified using two methods, peak-to-peak amplitude and area methods commonly used in other studies (Ellaway et al. 1998; Pascual-Leone et al. 1998; Thickbroom et al. 1999). MEPs in upper limb muscles were observed within the time period lasting from 10 ms to 50 ms after the TMS pulse (Fig. 2A). The peak-to-peak MEP amplitude was calculated by subtracting the minimum from the maximum of the EMG trace within this time period (Fig. 2B). The MEP area was calculated by first rectifying the EMG trace within this time period and then integrating it (Fig. 2C).

**Figure 2.**
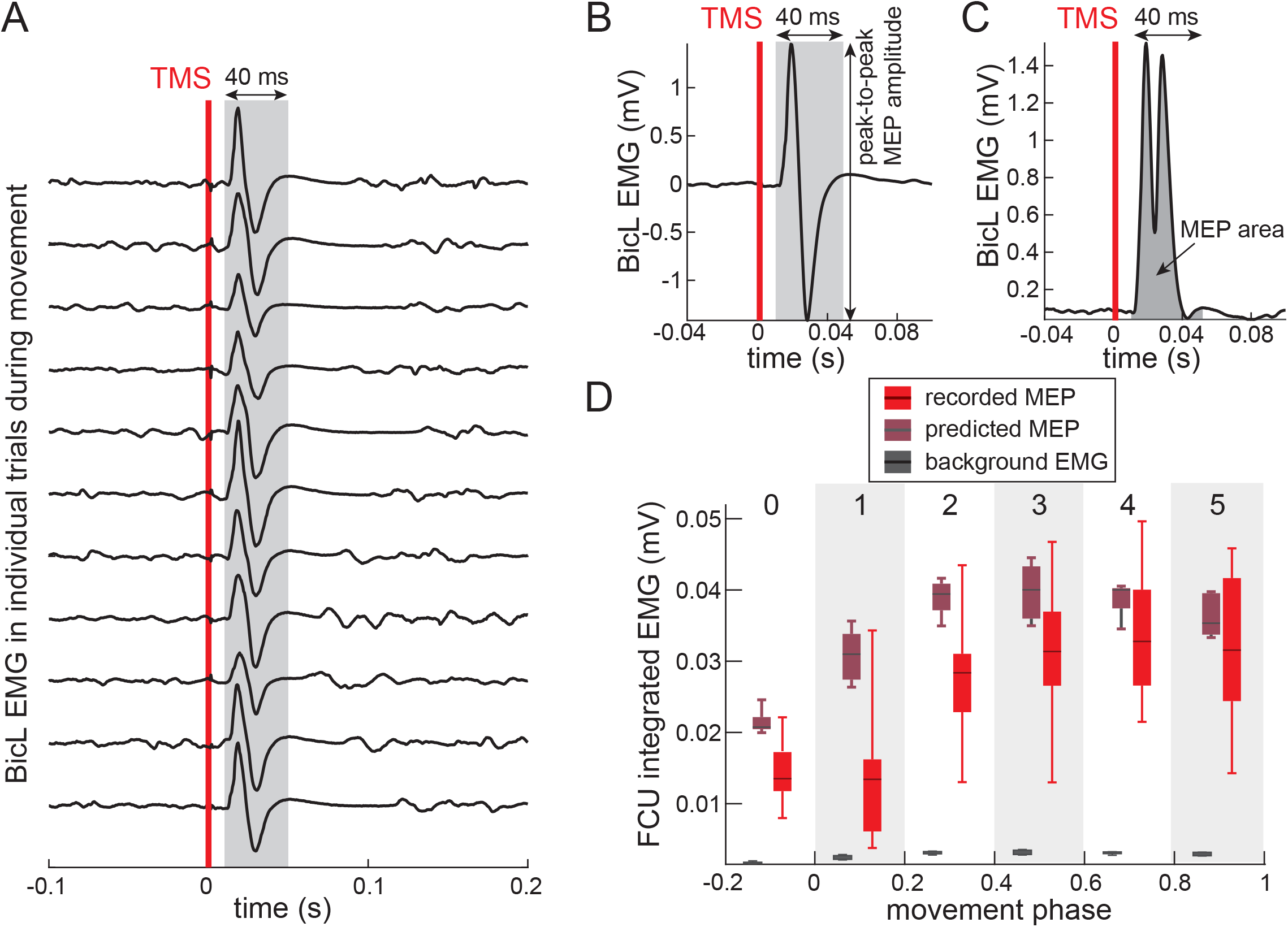
Quantifying MEP amplitude. A. Biceps EMG trace aligned on TMS pulse (red line) during reaching trials from the Control task in one individual. Grey area shows the window used to quantify MEP amplitude and MEP area. B. Averaged EMG from BicL muscle in one individual performing the Control task. EMG was aligned on TMS pulse (red line) and peak-to-peak MEP amplitude was calculated with the shaded area. C. Averaged rectified EMG from BicL muscle in one participant performing the Control task. EMG was aligned on TMS pulse (red line) and MEP area was calculated by integrating the shaded area under the curve. D. EMG from FCU muscle in one participant performing the Assistive task. MEP area values were binned according to the phase of movement, where 0 denotes the kinematic onset of movement and 1 denotes the kinematic offset. Background EMG is from trials without TMS; predicted MEPs were calculated using equation (3).

For MEPs collected in Session 1, termed static MEPs, both peak-to-peak and area methods were included into the analysis. The background EMG was calculated over the 40-ms time window directly preceding the TMS pulse using the corresponding peak-to-peak or integrated method respectively. We defined the presence of a MEP as a peak-to-peak amplitude of at least 5 standard deviations above background EMG amplitude. The probability of evoking MEP was calculated by dividing the number of detected MEPs by the number of stimuli under the same conditions. To determine the consistency of MEPs based on MEP area metric, we calculated the coefficient of variation (CV) across repetitions of the same task per individual per muscle.

The peak-to-peak MEP amplitude (*MEP*_*amp*_) during isometric contraction has been shown to follow the Boltzmann equation (Darling, 2006; Devanne, 2002):

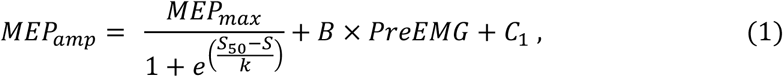

where *MEP*_*max*_ is the maximum MEP amplitude, S_50_ is the stimulation intensity to elicit a MEP at 50% of the maximum amplitude, S is the stimulation intensity, and *PreEMG* is the amplitide of EMG measured directly preceding stimulation, and *C*_1_ is a constant. In this study, we performed all stimulations at a consistent intensity of 90% the RMT, which makes the first term a constant. Therefore equation (1) can be simplified and adapted for examining the linear relationships between EMG and MEP area as follows:

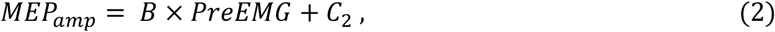

where *MEP*_*amp*_ in our experiment is MEP area measured as described above, *PreEMG* term is the background EMG calculated as describe above, and *B* is the slope of the linear relationship between MEP and EMG during posture maintenance. *C*_2_ was assumed to be equal to 0 expecting no MEPs in fully relaxed muscles due to stimulation below RMT. We derived the slope *B* in static Session 1 and used it to normalize the MEP area values recorded during movement in Session 2.

We calculated MEP latencies using a procedure similar to that used to determine kinematic onsets. Unrectified EMG signals were aligned on the TMS pulse and averaged across repetitions of the same postures. The MEP latency was defined as the time from the TMS pulse to the first maximum of the third derivative of this averaged trace.

For MEPs collected in Session 2, termed dynamic MEPs, both peak-to-peak and area methods were included into the analysis. All trials during which TMS was applied were included into the MEP metrics without determining directly whether a MEP was evoked or not in a given trial. This is because of difficulties distinguishing MEPs from compound motor unit action potentials in presence of changing EMG during movement. MEP area and the CV values were calculated as in Session 1. The MEP area values obtained from single trials were grouped into 5 temporal bins, each corresponded to 20% increments of phase duration from onset to offset of movement (Fig. 2D). This binning procedure grouped MEPs occurring at similar times during movement and provided adequate repetitions to estimate median values. Additionally, TMS responses that occurred up to 20% phase duration prior to movement were grouped into a bin 0 (Fig. 2D). We defined a minimum repetition criterion of 5 MEPs based upon (Lewis et al. 2014). Bins that contained < 5 MEPs were excluded from subsequent analyses. The MEP amplitude values were normalized by the median MEPs in the starting posture for the corresponding task and muscle from Session 1. Thus, MEP area values equal to 1 indicated that the MEPs during movement were equal to those at the starting posture for that movement.

To estimate the changed in corticospinal excitability that were independent from the motoneuronal excitability we calculated MEP gain. For this measure we sampled background EMG in trials without TMS at the same phase as TMS pulses in other trials of Session 2. We selected short chunks of EMG at the corresponding phase of movement as MEPs and processed them the same way as the MEP area or perk-to-peak metric (Fig. 2C). We then used the linear MEP-EMG relationship observed in static postures in Session 1 to predict MEP amplitude (and *MEP*_*Pred*_) for a given background EMG level using the following equation:

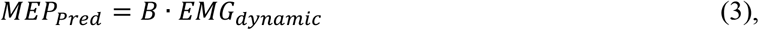

where *EMG*_*dynamic*_ is the median EMG within a given bin per muscle, task, and individual and slope *B* is from Eq. (2).

The MEP gain was calculated as the ratio between the observed MEP area (*MEP*_*Dyn*_) and *MEP*_*Pred*_ from Eq. (3). The MEP gain equal to 1 means that the median MEP at a given phase is equal to the predicted MEP based on the linear relationship between EMG and MEPs described in eq. (2) with the same slope as that observed in static trials. MEP gain values < 1 indicate a decrease in corticospinal contribution to a given muscle relative to that observed in static trials. By controlling for the linear relationship between EMG and MEP, the MEP gain metric reveals changes in the corticospinal tract that are independent of motoneuronal excitability.

Results in text are reported as mean values ± standard deviation across individuals, unless otherwise indicated.

### Statistical Analyses

The amount of shared variance between median MEP area values and the background EMG were calculated using the coefficient of determination (R^2^) across all movement phases per task per muscle per individual. For static MEPs, the background EMG was the value of *PreEMG* from Eq. (2). For dynamic MEPs, the background EMG was the value of *EMG*_*dynamic*_ from Eq. (3). The median MEP area was calculated as described above for each session. For data from Session 1, the linear regressions were fitted between MEPs, EMG, and the postural torques across all 5 postures (Fig. 1B). For data from Session 2, the linear regressions were fitted between MEPs and EMG across movement phases (bins 0 – 5). Significance was determined based on the goodness of fit statistic with Sidak correction for repeated testing setting α = 0.0043. The between-task differences in individual R^2^ values were normally distributed based on a single-sample Kolmogorov-Smirnov test (kstest function in MATLAB). Therefore, the significance of between-task differences was determined using paired t-tests on individual R^2^ across tasks using a ttest function in MATLAB. A separate test was done for each muscle using Sidak-corrected α = 0.017.

To determine if MEP gain was changing during movement, the values were divided into early and late groups based on whether MEPs occurred in the acceleration or deceleration phases of reach respectively. The early group comprised MEP gain values from bins 0, 1, and 2 and the late group comprised values from bins 4 and 5 (Fig. 2D). Bins 0 and 1 usually included fewer MEPs, therefore the total number of values in the two groups was comparable, > 40 in each acceleration or deceleration phase per muscle per individual per task. Single-trial MEP gain values were not normally distributed. Therefore, significance was determined using a two-sample Kolmogorov-Smirnov test using kstest2 function in MATLAB. Tests were applied to compare single-trial MEP gain values occurring in the acceleration and deceleration phases of movement for each individual per muscle with Sidak-corrected α = 0.0051. In contrast, the differences between median MEP gain values were normally distributed based on a single-sample Kolmogorov-Smirnov test. Therefore, paired t-tests were used to determine if MEP gain was modulated during movement per task per muscle with Sidak-corrected α = 0.017.

To quantify the changes in MEP gain between tasks, differences between individual median MEP gain values were calculated between tasks separately for acceleration and deceleration phases for each muscle. Then, corresponding differences in joint angle, muscle torque and its components were calculated for each individual per DOF. The joint angles in the acceleration (bins 0, 1, and 2) and deceleration (bins 5 and 6; Fig. 4A) phases of movement were averaged per individual per DOF and subtracted between tasks. The postural torques in the acceleration (bins 0, 1, and 2) and deceleration (bins 5 and 6; Fig. 4B and C) phases of movement were also averaged per individual per DOF and subtracted between tasks. For the dynamic component, maximal values in the acceleration phase and minimal values in the deceleration phase due to changing sign were calculated per individual per DOF and subtracted between tasks (Fig. 4D). The resulting difference values were normalized to their range across all individuals. Lastly, liner regressions were fitted separately for acceleration and deceleration phases of movement between each of the kinematic and dynamic variables and MEP gain of muscles that span the corresponding joint (Table 1). Sidak correction for repeated regressions was used to set α = 0.0127.

## Results

Our evidence supports the involvement of the corticospinal tract in posture maintenance in humans. At stimulation amplitudes below RMT, we observed MEPs during static posture, termed static MEPs, in most muscles. For example, biceps muscle participates directly in posture maintenance by holding the elbow and shoulder flexed in the tested postures (Fig. 3A). We found that the probability of evoking a MEP in this muscle was higher than expected from stimulation at the RMT in most individuals (Fig. 3B). Furthermore, the probability of evoking a static MEP was higher than expected in all muscles, even those not directly involved in supporting the arm against gravity (Fig. 3C). Altogether this shows that the overall corticospinal excitability was increased in a consistent manner when the arm was held against gravity in different postures compared to when the arm was relaxed.

**Figure 3.**
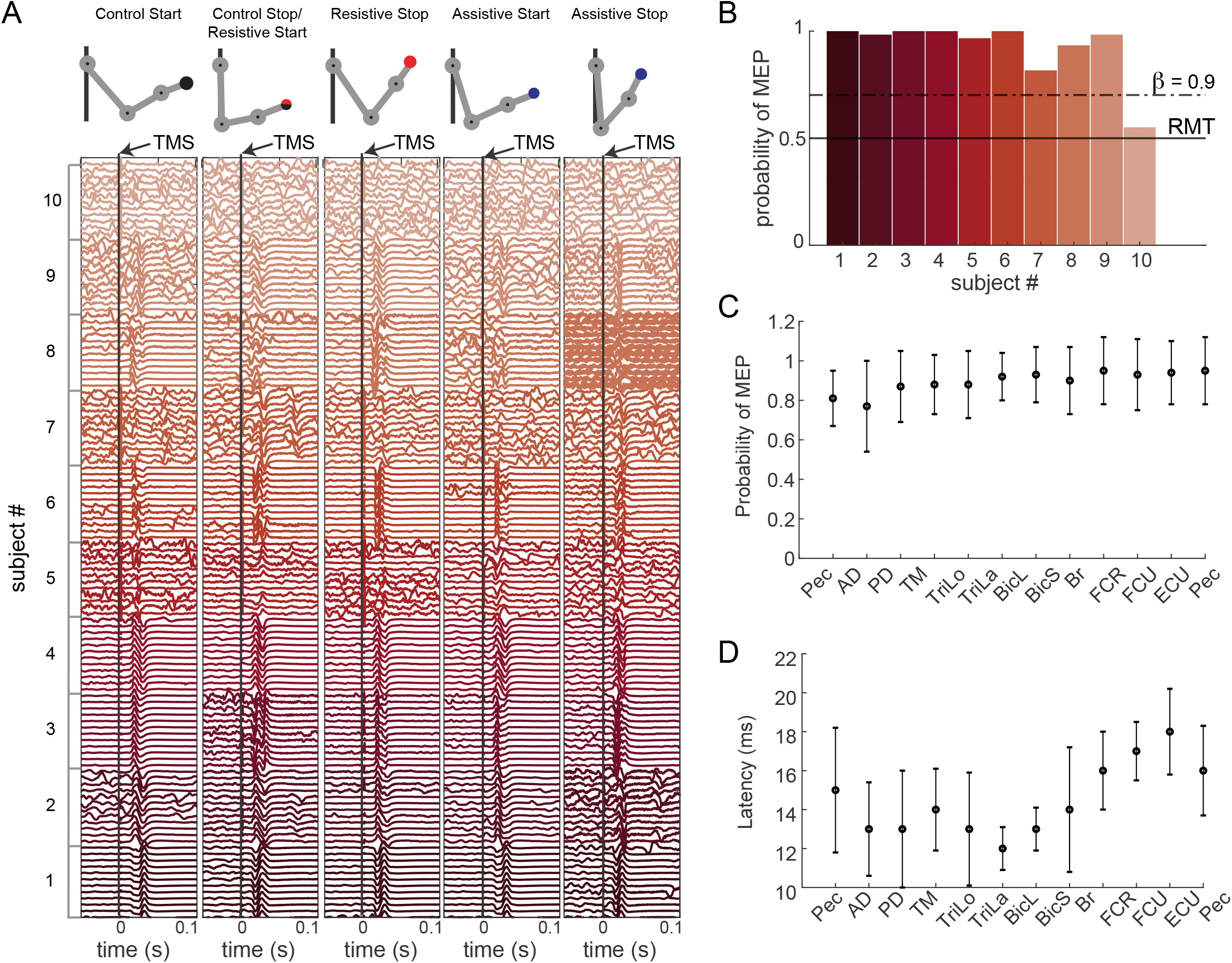
Static MEPs in BicL. A. EMG traces of the biceps muscle during static postures are displayed for all participants. The black line corresponds to the time of TMS pulse (stim). B. Each bar corresponds to the probability of a MEP occurring for each participant across all postures. The solid black line denotes the probability of MEP occurrence at RMT. The dashed line corresponds to the increase in probability that could be detected given the statistical power of our study design (β = 0.9).

The static MEPs were evoked at similar latencies and with consistent amplitudes across individuals. The average latency across muscles was 15 ± 1.9 ms (Fig. 3D). These latencies broadly reflected the proximal to distal distribution of muscles and the associated differences in the distance travelled by the action potentials evoked by TMS.

The MEP amplitudes were highly consistent as evidenced by the CV of static MEPs equal to 0.35 and dynamic MEPs equal to 0.5 (Fig. 4A), which is comparable to reported values recorded at rest and in relaxed muscles (Ellaway et al. 1998; Jung et al. 2010; Wassermann 2002; Zanette et al. 1995). The larger CV for the dynamic MEPs compared to the static MEPs is likely driven by the changing motoneuronal excitability reflected in the changing background EMG and muscle lengths during movement (Day et al. 1991; Kiers et al. 1993; Thickbroom et al. 1999). Furthermore, we observed that the static MEPs were linearly related to the changes in background EMG across postures (Fig. 4B). This linear relationship expressed in eq. (2) accounted for 25% of the static MEP variance (mean R^2^ = 25% ± 13% across muscles) with a small mean squared error of 2.2e^-5^ ± 4e^-5^. The regression slopes were above unity in all muscles and individuals (Table 1, static MEP / EMG column). These slopes also broadly reflected the proximal to distal distribution of muscles with distal muscles trending toward higher slopes than proximal ones. The dynamic MEPs displayed many similar characteristics to the static MEPs. Dynamic MEPs were also observed in all reaching tasks and in all muscles despite subthreshold stimulation. Dynamic MEP latencies were on average 14.2 ± 2.5 ms, similar to static MEPs. The dynamic MEPs were linearly related to the changes in background EMG during movement (Fig. 4C). In all muscles the ratios between dynamic MEPs and EMG in corresponding bins were above one and the ratios were similar or in some muscles above the static slopes (Table 1, dynamic MEP / EMG column).

**Figure 4.**
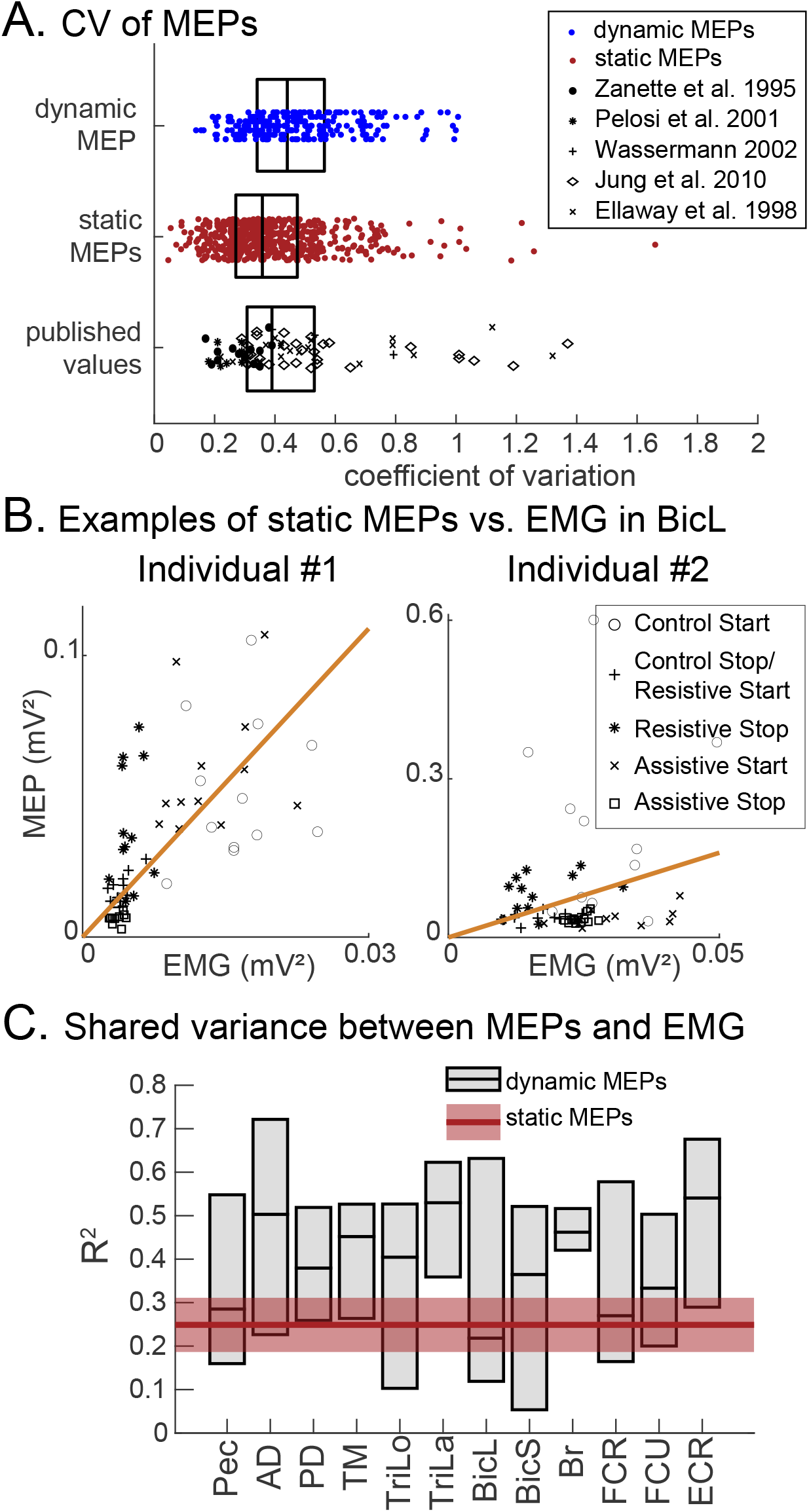
A. Coefficient of variation (CV) across all muscles and postures or movement per individual. CV for static MEPS are shown in red. CV from published studies are shown in black. CV for dynamic MEP gain values are shown in blue calculated as described in Methods, i.e. by dividing recorded MEP by predicted MEP from the corresponding bins (Fig. 2D). B. An example of the “best” (left) and “worst” (right) linear fit in BicL from different individuals. Lines are mean trajectories; error bars are standard deviations across individuals. C. Variance of dynamic MEPs explained by background EMG in individual muscles across tasks. Horizontal lines show medians and bars show quartile ranges across individuals. The horizontal red line and shaded area represent the mean variance of static MEPs explained by background EMG.

The virtual targets were effective in evoking the desired behavior and standardizing the movement kinematics and dynamics across all individuals (Fig. 5). Holding the arm against gravity in the five postures and transitioning between them during reaching was accomplished with postural torques of different magnitude at the major arm joints (Fig. 5B). The postural torques were larger during Session 1 (Fig. 5B, hold) than at the start and end of movement during Session 2 (Fig. 5B, bin 0 and 5) because of the additional contribution of the propulsive torques to accelerate and descelerate the arm during movement. However, the differences between postural torques across tasks were the same in both conditions. For example, the postural torque about the shoulder in the starting position for the Resistive condition was smaller than that in the Control and Assistive conditions in both Sessions (Fig. 5B, Shoulder). The shoulder motion in the Control and Assistive tasks was with gravity, both were accompanied by decreasing postural torque at the shoulder, while in the Resistive task the shoulder motion was against gravity and the postural torque was increasing (Fig. 5B Shoulder). In contrast, the postural torques about the elbow and wrist joints were more alike in the Assistive and Resistive (Fig. 5B, Elbow & Wrist, blue and red) tasks despite different motion of the elbow in these two tasks (Fig. 5A, Elbow & Wrist, blue and red). The postural torques at the elbow and wrist joints were equal across all starting postures and in the beginning of movement; they were diminishing during reaching in the Assistive and Resistive tasks but not in the Control task. These results suggest that the postural control of the elbow and wrist can be coupled, while the postural control of the shoulder needs to be decoupled from that of the elbow and wrist to appropriately compensate for the changing gravity-related passive torques. This applies to both posture-related component of focal movement and maintaining static posture.

**Figure 5.**
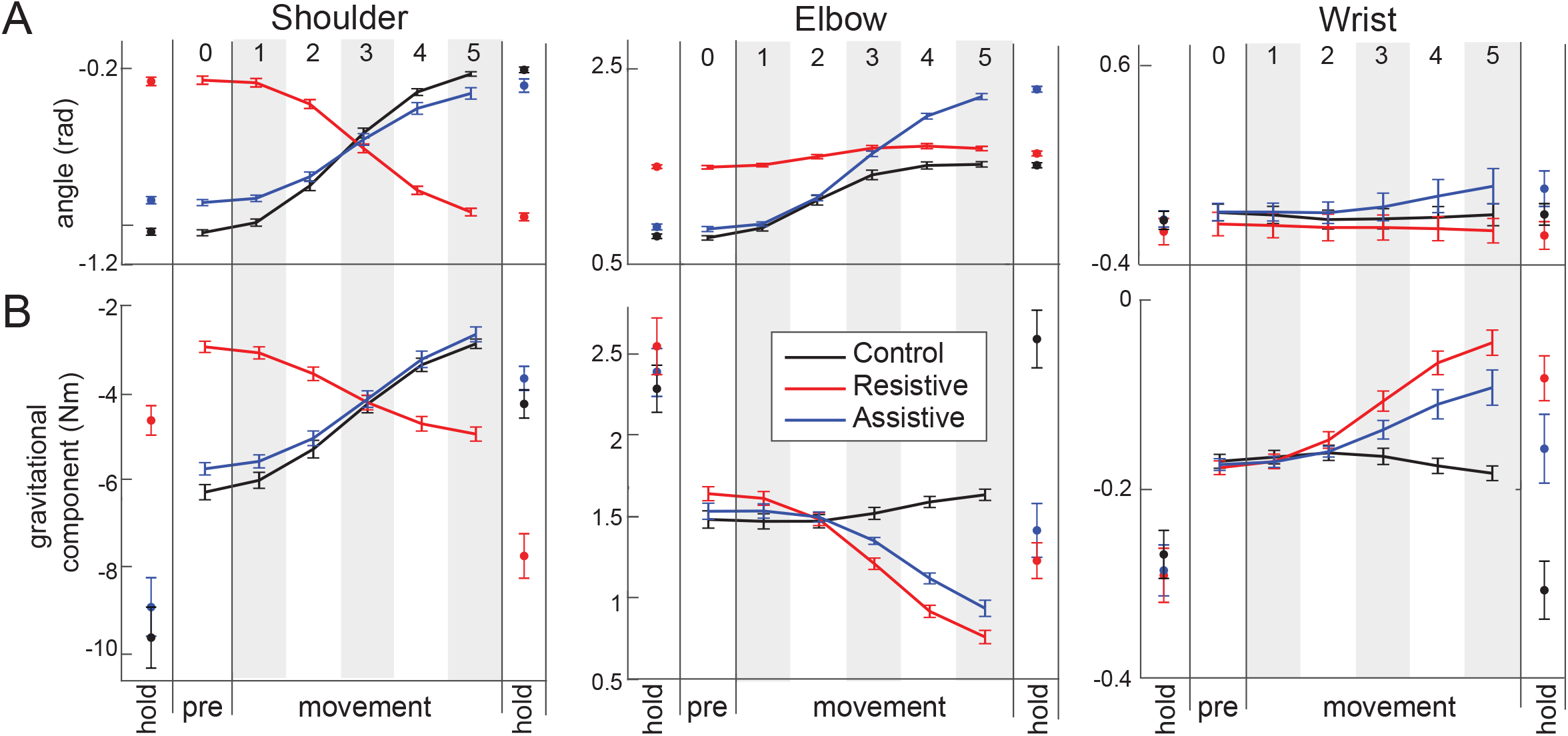
Movement kinematics and dynamics. A. Joint angles calculated from motion-capture B. The component of muscle torque that compensates for gravity load on the joints, termed postural torque.

The temporal profiles of postural torques during reaching can be used to define the tonic components of EMG in multiple muscles (Olesh et al. 2017). Here too we observed that in Session 1 the EMG amplitude in most muscles spanning the shoulder was linearly dependent on the amplitude of shoulder postural torque across the five postures with shared variance for Pec = 30%, AD = 65%, PD = 38%, TM = 31%, TriLo = 45% at p < 0.001 for each (significant α = 0.004). In contrast, the EMG amplitude of muscles spanning the elbow or wrist was weakly dependent on the amplitude of elbow or wrist postural torques (R^2^ for TriLo = 6%, TriLa = 4%, BicL = 1%, BicS = 6%, FCR = 2%, FCU = 7%, ECR = 5% at p > 0.07 for each and BicL = 11% at p = 0.017, significant α = 0.004). Furthermore, the static MEPs only in the muscles spanning the shoulder were linearly related to the shoulder postural torque in Session 1 with high shared variance (R^2^ for Pec = 49%, AD = 51%, PD = 43%, TM = 47%, TriLo = 48% at p < 0.001 for each; significant α = 0.004). Interestingly, removing the static MEP dependency on the background EMG by calculating static MEP gain did not fully remove the MEP dependency on the postural torque. There remained a linear dependency between shoulder postural torque and static MEP gain in Pec (R^2^ = 19% at p = 0.002) and in TriLo (R^2^ = 23% at p < 0.001), indicating modulated descending contribution to the recruitment of these muscles. In the rest of the muscles, the static MEPs were not correlated with the changes in the elbow or wrist postural torques across the 5 static postures. This shows that to hold the arm against gravity, the descending contribution to most muscles was elevated. Furthermore, the descending contribution to the recruitment of Pec and TriLo was modulated with the shoulder postural torque.

The dynamic MEP gain values during movement varied more between individuals than between muscles. This suggests that the inter-subject variability may contain biomechanically relevant information rather than noise. To test this idea, we performed a regression analysis, in which we included only the dynamic MEP gain values per individual per muscle that were significantly different between tasks in a corresponding phase of movement. The MEP gain in monoarticular shoulder muscles (Pec, AD, PD, and TM) was not correlated with the changes in shoulder postural torque at the beginning of movement (Fig. 6A). This shows that change in the descending contribution to the recruitment of monoarticular shoulder muscles does not serve the role of counteracting the passive postural torque at the shoulder in the beginning of movement. In contrast, the dynamic MEP gain in the biarticular biceps and triceps muscles (BicL and TriLo) was increased when the arm was more outstretched, i.e. at larger shoulder flexion angle and lower elbow flexion angle (Fig. 5, black and blue). The outstretched postures in Assistive and Control tasks were accompanied by larger shoulder postural torque, while the elbow torque remained unchanged. Consequently, the dynamic MEP gain in BicL and TriLo was linearly proportional to the increase in shoulder postural torque between tasks (Fig. 6B). The elbow postural torque was similar between tasks in the beginning of movement (Fig. 5B), therefore no correlations with MEP gain in the muscles spanning the elbow were observed (Fig. 6C). The wrist angle and postural torque were similar across tasks in the beginning of movement (Fig. 5) and thus no correlations with MEP gain in wrist muscles (FCU, FCR, and ECR) were observed (Fig. 6D). This shows that the inter-subject variability of the dynamic MEP gain in BicL and TriLo was linearly dependent on the individual differences in the postural torques in the beginning of reaching tasks. Furthermore, these results suggest that the changes in the descending contribution to the recruitment of biarticular biceps and triceps muscles in the beginning of movement compensate for the interaction torques between shoulder and elbow caused by the changing passive gravity torques at these joints.

**Figure 6.**
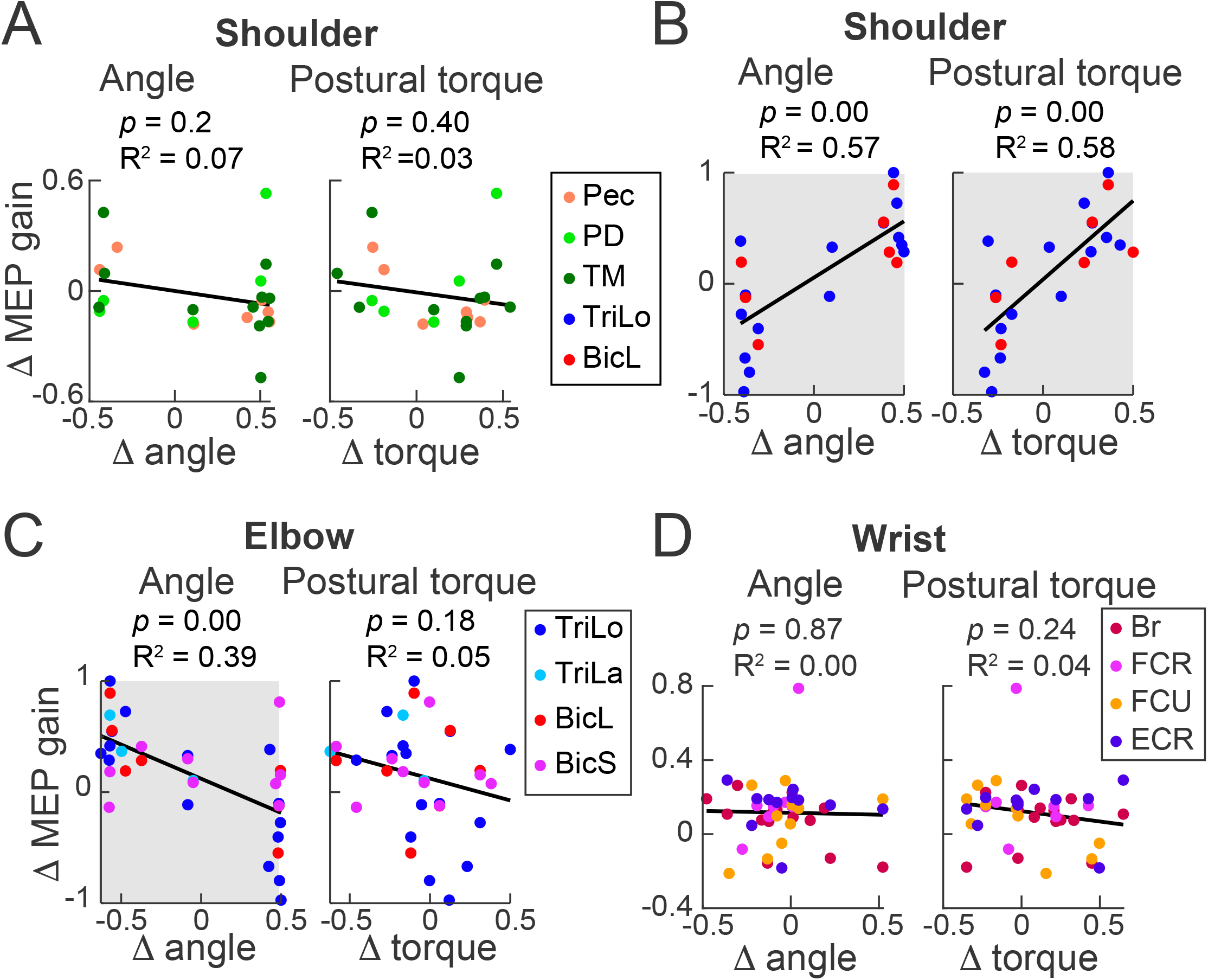
Modulation of MEP gain between tasks in the beginning of movement. Circles show MEP gain differences between tasks per individual per muscle. Thick black lines show regressions fitted to all values in the plot with goodness of fit statistic (*p* and R^2^) above each plot. Plots with significant linear dependencies at alpha < 0.0127 have grey background. A. Differences in MEP gain between tasks in pectoralis (Pec), posterior deltoid (PD), and teres major (TM) relative to the differences between tasks in shoulder angle and in shoulder postural torque. B. Differences in MEP gain between tasks in triceps (long head, TriLo) and biceps (long head, BicL) relative to the differences between tasks in shoulder angle and shoulder postural torque. C. Differences in MEP gain between tasks in triceps (long and lateral heads, TriLo and TriLa respectively) and biceps (long and should heads, BicL and BicS respectively) relative to the differences between tasks in elbow angle and elbow postural torque. D. Differences in MEP gain between tasks in brachialis (Br), flexor carpi ulnaris (FCU), flexor carpi radialis (FCR), and extensor carpi radialis (ECR) relative to the differences between tasks in wrist angle and wrist postural torque.

At the end of movement, the MEP gain in the monoarticular shoulder muscles was proportional to the shoulder and wrist postural torques respectively (Fig. 7A). In contrast, the dynamic MEP gain in the biceps and triceps muscles did not depend on either shoulder or elbow postural torque (Fig. 7B, 7C) despite large differences in these torques between tasks (Fig. 5). Furthermore, the MEP gain in the wrist muscles and brachioradialis was proportional to the wrist postural torque in absence of motion around the wrist (Fig. 7D). These results suggest that the postural adjustments to stabilize the arm at the new posture at the end of movement are driven by modulated descending contribution to the recruitment of multiple monoarticular shoulder muscles, brachioradialis, and all recorded muscles spanning the wrist. The changes in the descending contribution to proximal muscles serve the role of counteracting the passive postural torque at the shoulder, while changes in the descending contribution to distal muscles, including brachioradialis, serve the role of counteracting the passive postural torque at the wrist. However, the descending contribution to the recruitment of biceps and triceps muscles at the end of movement does not serve the role of counteracting the passive postural torque at the joints that these muscles span.

**Figure 7.**
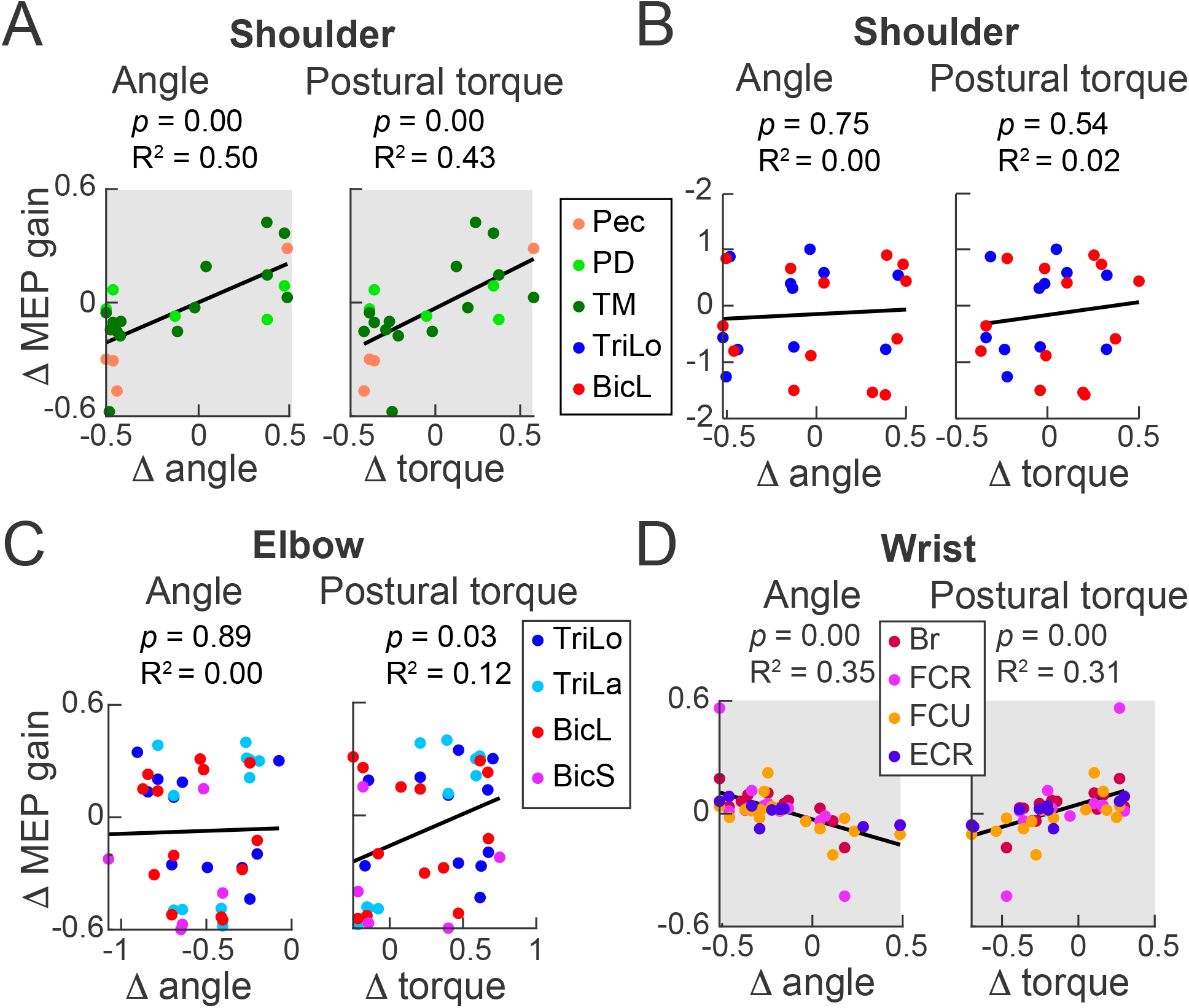
Modulation of MEP gain between tasks at the end of movement. Circles show MEP gain differences between tasks per individual per muscle. Thick black lines show regressions fitted to all values in the plot with goodness of fit statistic (*p* and R^2^) above each plot. Plots with significant linear dependencies at alpha < 0.0127 have grey background. A. Differences in MEP gain between tasks in pectoralis (Pec), posterior deltoid (PD), and teres major (TM) relative to the differences between tasks in shoulder angle and in shoulder postural torque. B. Differences in MEP gain between tasks in triceps (long head, TriLo) and biceps (long head, BicL) relative to the differences between tasks in shoulder angle and shoulder postural torque. C. Differences in MEP gain between tasks in triceps (long and lateral heads, TriLo and TriLa respectively) and biceps (long and should heads, BicL and BicS respectively) relative to the differences between tasks in elbow angle and elbow postural torque. D. Differences in MEP gain between tasks in brachialis (Br), flexor carpi ulnaris (FCU), flexor carpi radialis (FCR), and extensor carpi radialis (ECR) relative to the differences between tasks in wrist angle and wrist postural torque.

## Discussion

Here we used single-pulse TMS to probe corticospinal excitability during unconstrained postural and reaching tasks. We observed that stimulation below RMT evoked MEPs in all muscles during both holding arm in static posture and during reaching. This provides evidence that both maintaining a static posture against gravity and transitioning from one posture to another involve increased motorcortical contribution to the recruitment of most muscles of the arm. Under all conditions MEPs were linearly dependent on the background EMG with 25% of variance in static MEP amplitude and even more in dynamic MEP amplitude captured by this linear relationship (Fig. 4C). This is consistent with previous work showing a large amount of MEP variance is captured by a simple linear relationship between MEP amplitude and EMG during isometric voluntary contractions (Darling et al. 2006). The slope of this linear relationship quantifies a certain level of the motorcortical contribution to the recruitment of the corresponding muscle. Any changes in this slope or changes in MEP gain reflect the change in the level of motorcortical contribution to the recruitment of a given motoneuron pool. For example, the slopes of the linear relationship between MEP and EMG in distal muscles tended to be higher than proximal muscles (Table 1). This is consistent with prior work showing a greater involvement of the corticospinal tract in the voluntary activation of distal vs. proximal muscles (Rothwell et al. 1991; Turton and Lemon 1999). In our study the MEP gain in pectoralis and the biarticular triceps muscles changed in proportion to the shoulder postural torque during posture maintenance. Moreover, in the beginning of movement, the MEP gain in both biarticular biceps and triceps muscles changed in proportion to shoulder postural torque. At the end of movement, the MEP gain in the monoarticular shoulder muscles (pectoralis, posterior deltoid, and teres major) was proportional to the shoulder postural torque and the MEP gain in wrist muscles (flexor carpi radialis, flexor carpi ulnaris, and extensor carpi radialis) was proportional to the wrist postural torque. This shows that the motorcortical contribution to the recruitment of more arm muscles increased in proportion to postural torques during movement compared to that during posture maintenance. Altogether, our results support the hypothesis that the motor cortex contributes to muscle recruitment to both maintain posture and to compensate for changes in gravitational passive joint moments during movement.

The reaching tasks used in our study were selected to maximize the differences in the postural forces that occur at the major arm joints while minimizing the differences in elbow and wrist kinematics. Consequently, large differences in the postural components of muscle torques were needed to hold the arm in different postures or to transition between postures (Fig. 5). The transitions between postures required modulated motorcortical involvement in the recruitment of muscles forming distinct groups. In the acceleration phase of movement, the MEP gain in both antagonistic biarticular biceps and triceps muscles was proportional to the shoulder postural torque (Fig. 6B). This suggests that at the beginning of movement the motorcortical projections contribute to the coactivation of biceps and triceps muscles (Fig. 8, red projections). This may serve the purpose of compensating for the interaction torques caused by the changing gravity load at the shoulder through increasing mechanical impedance of shoulder and elbow joints (Gribble and Ostry 1999; Gritsenko et al. 2011). This was followed by the MEP gains in proximal and distal muscles being proportional to the shoulder and wrist postural torque respectively at the end of movement (Fig. 7A, 7D). These results are consistent with earlier observations of phase-specific modulation of corticospinal gain in multiple arm and hand muscles during reach and grasp movement (Lemon et al. 1995, 1996). Our results indicate more specifically that in the deceleration phase of movement the motorcortical projections contribute to the coactivation of subsets of proximal and distal muscles, which likely increased the mechanical impedance of shoulder and wrist joints and stabilized the arm in the new posture (Fig. 8, blue projections). We postulate that the postural torque in both acceleration and deceleration phases of movement reflect postural adjustments that accompany focal movement, as it is precisely the active joint torque needed to support the arm against gravity. Therefore, the observed modulation of MEP gain with the postural torque both in our postural and reaching tasks is likely to involve the reticulospinal tract via collaterals from the corticospinal tract (Keizer and Kuypers 1989). The importance of the reticulospinal tract in the integration between posture and movement, including reach, is well established in cats (Schepens and Drew 2004, 2006) and primates (Baker 2011; Riddle et al. 2009). The TMS methodology used here may help study the maladaptive changes in the neural mechanisms of postural adjustments in humans with neurological conditions and identify avenues of new neuromodulative interventions.

**Figure 8.**
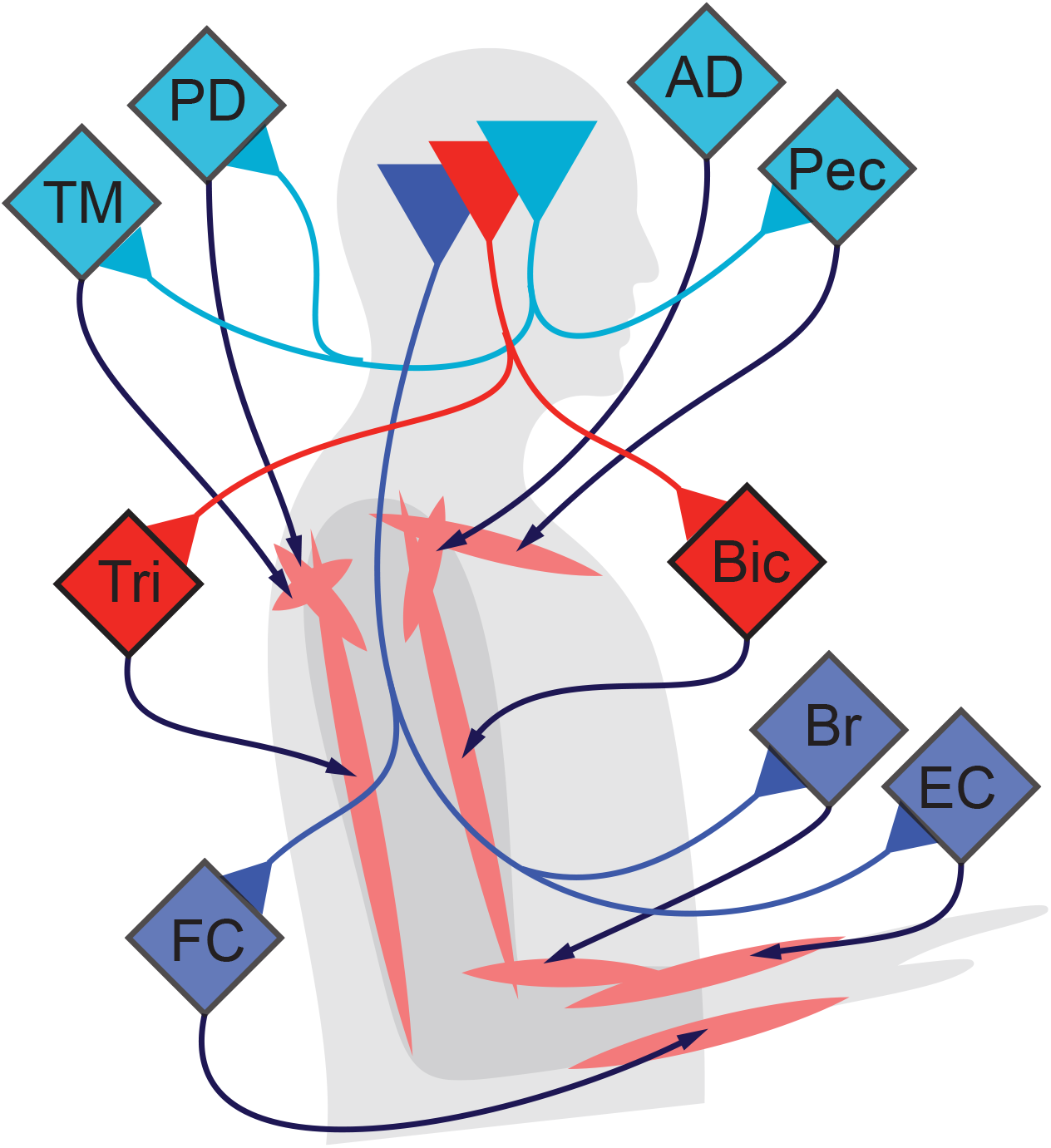
Concept of the control schema for postural adjustments during reaching. A. Motorcortical outputs for counteracting the changing gravity load on the arm. Grey outlines the human body and arm; spindles show the basic anatomy of major muscle groups. Diamonds indicate motoneuron pools of the corresponding muscles. They are colored to indicate common input from the pyramidal cells in the primary motor cortex (triangles). Colors of the pyramidal contributions indicate changes in MEP gain in the acceleration phase of movement (red) and in the deceleration phase of movement (blue).

The limitations of study methodology may limit the generality of our conclusions. The first limitation is that the contribution of descending projections from premotor and supplementary motor cortices to motoneuronal excitability cannot be ruled out. In our study we used focal TMS that was targeted and scaled based on responses observed in biceps EMG. This method is designed to limit the volume of current density that is sufficient to stimulate cells located in the arm area of the primary motor and possibly primary sensory motor cortices (Pascual-Leone et al. 1997; Pascual-Leone and Walsh 2002). Therefore, the descending projections from premotor areas were likely not stimulated and their contribution to muscle recruitment and the compensation for passive torques not quantified in the present study. The second limitation is the possibility that using a different muscle for determining the RMT would alter the modulation of MEP gain in other muscles. It seems unlikely for proximal muscles, because the spatial resolution of TMS is rather low. Stimulation over scalp locations 1-2 cm apart has been shown to crudely distinguish between proximal or distal muscle groups (Pascual-Leone et al. 1997). Here we have targeted a proximal muscle group by determining our hot-spot and RMT based on MEPs in biceps. If we choose another proximal muscle for thresholding, the hot-spot method will likely zero in on the same scalp location but a different magnetic field strength. Increasing or decreasing the magnetic field strength over the same spatial location in order to target another proximal muscle is likely to activate more or less respectively all the same muscles recorded here. The underlying cortical activity that drives the modulation of MEP gain would be the same. However, if we choose a distal muscle for thresholding, the hot-spot method will likely zero in on a different scalp location and a different magnetic field strength. In this case, proximal muscles are not likely to have MEPs due to their generally higher threshold (Rothwell et al. 1991; Turton and Lemon 1999). Therefore, the activity of the different underlying cortical area may cause different MEP amplitudes and gains across all muscles compared to those caused by the scalp area chosen based on the biceps hot-spot. Consequently, our results generalize to represent a subset of cortical contribution to the control of reaching, but they may not generalize to the more dexterous corticospinal control of distal muscles. The third limitation is that the MEPs produced by TMS during movement may saturate in some muscles at some phases of movement. This possibility is based on observations of similar or lower MEP amplitudes despite increasing background EMG from 10% to 40% of maximal voluntary effort for the former or 25% to >50% of maximal voluntary effort for the latter (Devanne et al. 1997; Kamen 2004). Here, we did observe a linear relationship between background EMG and MEP amplitude with similar slopes during both postural and reaching tasks (Table 1). Importantly, during reaching tasks the range of EMG changes is larger than during the different postures, yet we did not observe signs of saturation that would be evident from reduced slope of the regression during reaching tasks. However, the slopes for distal muscles (Br, FCR, FCU, and ECR) were rather high, which suggests a very high recruitment of these projections by TMS. Therefore, the potential saturation of corticospinal contribution to the recruitment of distal muscles probed with TMS in our study cannot be completely ruled out.

## Endnote

All data relevant for assessing the conclusions of this study are presented in the manuscript and figures.

## Acknowledgements

We thank the study participants for generously giving their time; we acknowledge the data collection and analyses contributions of Dr. W. Talkington; we acknowledge the technical support expertly provided by B. Pollard; we are thankful for insightful and critical comments of Dr. S. Yakovenko. The current affiliation of RLH is Albany Stratton VA Medical Center, Albany, NY.

## Figure Legends

**Extended Data 1**. Software and data are included in two folders and described in the readme.txt file. The code for inverse kinematic and dynamic simulations is in the Simulation Functions folder. To obtain joint angles from a given set of marker coordinates, run getEuler_ARM_140205.m. To obtain joint torques from a given set of joint angles, run MODEL_getTorques.m. This function runs a Simulink model of the arm KINARM_Dyn_Inv.mdl. The code for plotting figures and statistics is in the Manuscript tests and plots folder. Run plot_RSquaredAnalysis.m to plot figures 6 C and D from data saved in mat files. Run getDynamicMEP_plot_EarlyLateTest.m to plot figures 7, 8, and 9 with associated statistics using data saved in mat files. This script also contains a power test for one of the regressions.

